# Goal Uncertainty Attenuates Sensorimotor Adaptation

**DOI:** 10.1101/2025.07.18.665537

**Authors:** Sritej Padmanabhan, Reza Shadmehr, Roberta Klatzky, Jonathan S. Tsay

**Affiliations:** Department of Psychology, Carnegie Mellon University; Neuroscience Institute, Carnegie Mellon University; Department of Biomedical Engineering, Johns Hopkins University

## Abstract

Implicit sensorimotor adaptation—the automatic correction of movement errors through feedback and practice—is driven by a perceptual prediction error, the mismatch between the perceived movement outcome and its intended goal. While perceptual uncertainty is known to attenuate adaptation, the impact of goal uncertainty on adaptation remains unknown. We employed a visuomotor adaptation task that isolates implicit adaptation (N = 180), manipulating goal uncertainty by varying how precisely the goal’s midpoint could be identified. Display format was varied independently to control for the objective size of visual features, and targets were hidden at movement onset, ensuring identical visual input at the moment the error was experienced. We found that goal uncertainty significantly attenuated implicit adaptation, independent of low-level visual and kinematic features. Together, these results demonstrate that a precise internal representation of the goal is essential for supporting implicit sensorimotor adaptation.

## Introduction

Multiple learning processes support goal-directed action, enabling individuals to adapt to changes in their bodies (e.g., muscle fatigue) and environments (e.g., using a new ping-pong paddle) (1–5). Among these, implicit adaptation—the automatic correction of movement errors through feedback and practice—is essential for keeping the sensorimotor system well-calibrated (6,7). This form of learning is driven by perceptual prediction error: a mismatch between the perceived outcome of the movement and its intended goal (6,8–10). For example, if you reach for a cup and perceive your hand as landing rightward of it, the nervous system detects this error and updates future movements to improve accuracy.

Extensive prior work has shown that *perceptual feedback uncertainty*—noise in sensory signals about the movement outcome—attenuates implicit adaptation. For example, adaptation is reduced when visual feedback about a movement’s endpoint is spatially degraded—such as when it appears as a cloud of dots or is optically blurred (8,11–14). Similarly, individuals with blurring due to low vision exhibit diminished adaptation compared to those with normal vision (15). These effects are often explained by Bayesian principles: As feedback uncertainty increases, the perceptual prediction error becomes less reliable, leading the nervous system to downweight its influence—thereby attenuating implicit adaptation (16–19).

While uncertainty in spatial feedback is known to attenuate motor adaptation, a critical factor remains understudied: spatial uncertainty about the goal itself. The latter can arise either when sensory signaling of the goal lacks spatial precision, or when a goal signal that may be perceptually clear has low information value. As an analogy, a person giving directions may mumble or speak clearly but vaguely. As will be detailed in the discussion, while formal models of adaptive learning have focused on how feedback imprecision degrades the utility of the error signal, similar effects can be expected from imprecision in initial goal specification.

The neglect of goal uncertainty as a factor in motor adaptation is surprising, as the motor system is inherently goal-directed. Common neural circuitry has been implicated in the selection of an action goal and its execution (20), and emerging evidence shows that goal-related signals feed into the cerebellar circuits underlying implicit adaptation (21), suggesting that degraded goal representations could blunt learning at its source. Probing the effects of goal uncertainty on motor adaptation thus offers a principled test of how internal goal representations modulate cerebellar-dependent learning. Removing the visual target entirely is one test that might be proposed, but this manipulation introduces a confound between elimination of the task goal and changes in the low-level visual features that denote the target (e.g., edges and filled regions), and the effects of target removal have been inconsistent when tested (22–24).

To address these limitations, we used a visuomotor adaptation task that induces implicit adaptation through an irrelevant feedback signal (25,26). Our design manipulated the spatial precision of a designated target, while controlling for its low-level visual content and clearly defining the task goal: Participants made direct reaching movements to “slice through the midpoint of the target quickly and accurately,” while being instructed to “ignore the visual feedback,” which followed at a fixed spatial offset from the target, creating a consistent, task-irrelevant perceptual error. These instructions specified an unambiguous target location and removed the need for explicit re-aiming, ensuring that any deviation from the target reflected solely implicit adaptation.

Critically, we implemented a 2×2 design (N = 180) that crossed goal uncertainty with the size of visual display features. Goal uncertainty was manipulated by the spatial precision with which the target’s midpoint was specified: The low-uncertainty target was centered within a narrow region, while the high-uncertainty target was centered along a broad arc, which is more susceptible to bisection errors (27– 29). To dissociate goal uncertainty from visual extent, both high- and low-uncertainty targets were designated by small or large visual features. The target disappeared at movement onset, ensuring identical visual input across conditions at the moment of error feedback. If goal uncertainty attenuates implicit adaptation in a Bayes-optimal manner, targets specified by an arc should produce less adaptation than targets constrained to a small region, independent of the size of the features in the display.

## Results

How does goal uncertainty impact implicit motor adaptation? To address this question, we conducted a well-powered online visuomotor rotation study using a 2×2 between-subjects design. Participants (N = 180) were randomly assigned to one of four groups that varied in goal uncertainty (small vs large target) (27–29) and visual feature size (small vs large display) (Figure 1C). All participants were instructed to “slice through the midpoint of the target quickly and accurately.” The target disappeared at movement onset, serving as a go-cue and ensuring matched visual input at the time of error feedback.

**Figure 1.**
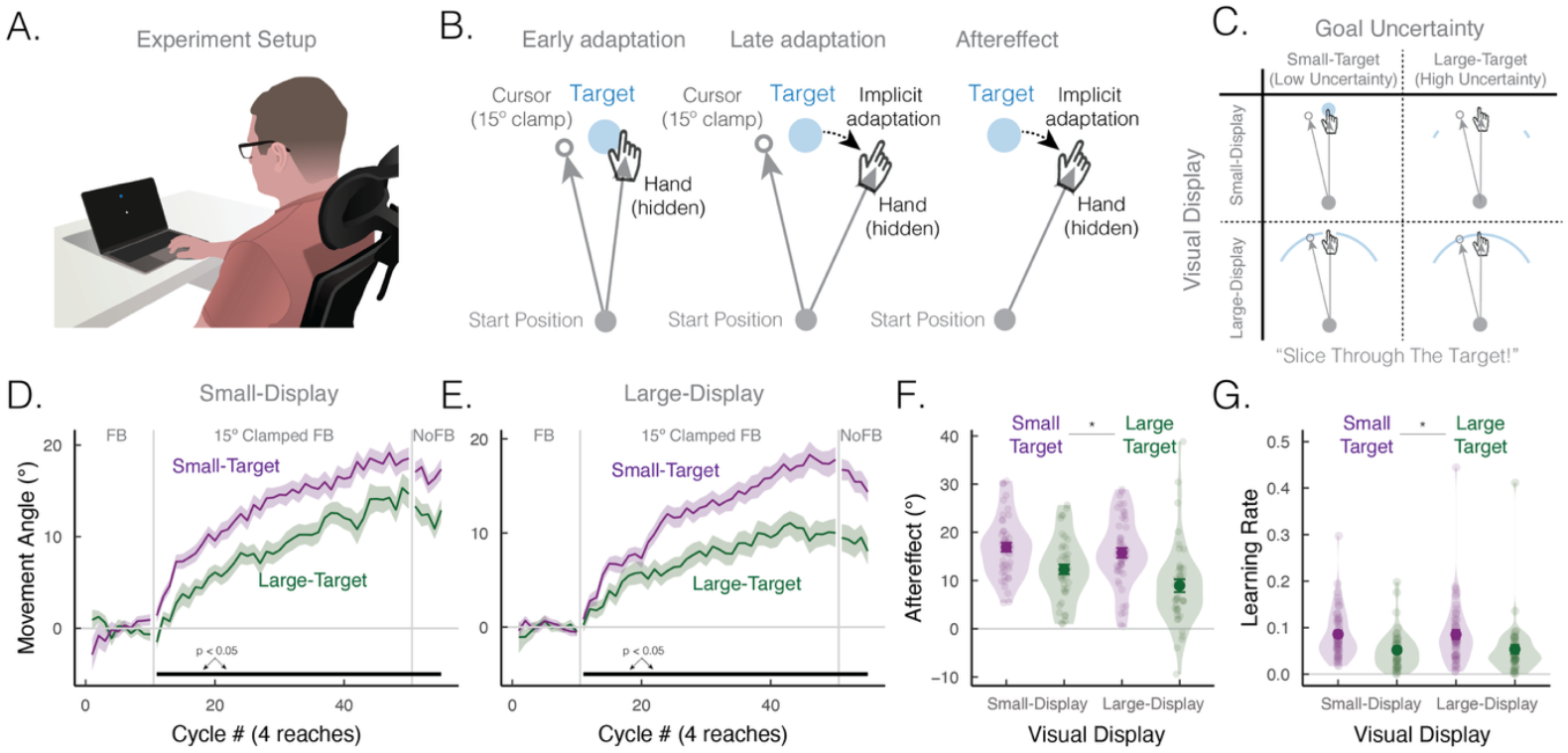
Goal Uncertainty Attenuates Implicit Sensorimotor Adaptation. **(A)** Experiment Setup. Participants used a trackpad to make rapid, goal-directed movements toward a visual target displayed on a laptop screen. **(B)** Clamped Feedback Task isolates Implicit Adaptation. After baseline trials with veridical cursor feedback (cycles 1–10), participants completed 160 trials with clamped feedback (cycles 11–50), in which the visual cursor followed a fixed trajectory rotated 15° clockwise or counterclockwise (counterbalanced across participants), regardless of actual movement direction. Participants were instructed to always “slice through the target’s midpoint” (blue circle) and ignore the cursor (white circle). Schematics illustrate cursor and hand positions during early adaptation (left: cycles 11–20), late adaptation (middle: cycles 41– 50), and aftereffects (right: cycles 51–55). Note that the target was removed during movement and at the time of feedback to ensure identical visual displays across groups. **(C)** Experimental Design. Participants were assigned to one of four groups in a 2×2 design manipulating the visual target (Small-Target vs. Large-Target) and visual display (Small-Display vs. Large-Display). High goal uncertainty (Large-Target) was induced using a spatially-extended arc lacking a clear midpoint; low goal uncertainty (Small-Target) used a small, delimited region for the center. **(D–E)** Mean Time Course of Implicit Adaptation. Participants exhibited greater implicit adaptation in Low-uncertainty conditions (purple) than High-uncertainty conditions (green), for both small **(D)** and large **(E)** displays. **(F-G)** Summary of Aftereffects and Learning Rate. Both implicit aftereffects **(F)** and learning rates derived from a standard state-space model **(G)** were reduced in the High-uncertainty groups compared to the Low-uncertainty groups. Dots represent individual participants; error bars indicate ±1 SEM around the mean. Asterisks (^*^) denotes p < 0.05.

After the baseline familiarization phase, participants were exposed to clamped visual feedback—a fixed 15° rotation independent of actual hand movement—to isolate implicit adaptation. Adaptation was measured by changes in movement angle across the perturbation and no-feedback aftereffect blocks. Despite being explicitly told the feedback was noncontingent and instructed to ignore it, all groups gradually shifted their movements opposite to the direction of the cursor (Figure 1D-E). This deviation persisted into the no-feedback block, confirming that the adaptation was driven by implicit processes.

To quantify adaptation, we first analyzed aftereffect performance (Figure 1E). All groups showed significant aftereffects compared to baseline (Low-Uncertainty /Small-Display: 16.9 ± 0.9°, Low-Uncertainty / Large-Display: 15.8 ± 0.8°, High-Uncertainty / Small-Display: 12.3 ± 0.9°, High-Uncertainty / Large-Display: 8.9 ± 0.9°; all p < 0.001). Critically, aftereffects were significantly reduced in the High-Uncertainty groups compared to the Low-Uncertainty groups (F(1, 176) = 6.6, p = 0.011). This effect was consistent across visual displays (main effect: F(1, 176) = 0.02, p = 0.90; interaction: F(1, 176) = 0.01, p = 0.91), indicating that the attenuation was driven by goal uncertainty rather than visual format.

A cluster-based permutation test reinforced this result, revealing a sustained reduction in adaptation in the High-Uncertainty groups throughout the perturbation and washout phases (cycles 11–55, F_sum_ = 832.5, p_perm_ < 0.001). A brief effect of visual display appeared only in the final two cycles (cycles 49– 50, F_sum_ = 10.0, p_perm_ = 0.02), with slightly greater adaptation for Small-Displays. However, this transient effect further suggests that display format had minimal influence on learning. No interaction between goal uncertainty and visual display emerged at any point.

To assess how goal uncertainty affects learning dynamics, we fit a state-space model to trial-by-trial movement angles (Methods). While retention rates did not differ significantly across groups (goal uncertainty: F(1, 176) = 2.8, p = 0.09; display: F(1, 176) = 1.1, p = 0.29; interaction: F(1, 176) = 0.2, p = 0.64), learning rates were significantly lower in High-Uncertainty groups (Figure 1G) (goal uncertainty: F(1, 176) = 6.6, p = 0.011), with no effect of visual display (F(1, 176) = 0.02, p = 0.90) and no interaction (F(1, 176) = 0.01, p = 0.91). Together, these results show that goal uncertainty slows both the rate and magnitude of implicit adaptation—independent of visual features.

An alternative explanation is that goal uncertainty may have influenced baseline motor control. For example, greater uncertainty could impact motor bias (mean movement angle), motor noise (standard deviation of movement angles), movement time (time from movement initiation at 1 cm to termination at target distance), or motor planning (e.g., reaction time, defined as the interval between target disappearance and movement initiation at 1 cm), thereby indirectly impacting adaptation (30,31). Importantly, goal uncertainty had no effect on kinematic measures (motor bias: t(177) = -0.5, p = 0.62; motor noise: t(177) = -0.08, p = 0.93; movement time: t(177) = 0.9, p = 0.34; reaction time: t(177) = 0.4, p = 0.68) (Figure 2). Together, these results indicate that the attenuating effect of goal uncertainty on adaptation cannot be explained by differences in motor execution, movement planning, or visual features. Rather, it reflects a fundamental consequence of increased uncertainty in the internal representation of the goal.

**Figure 2.**
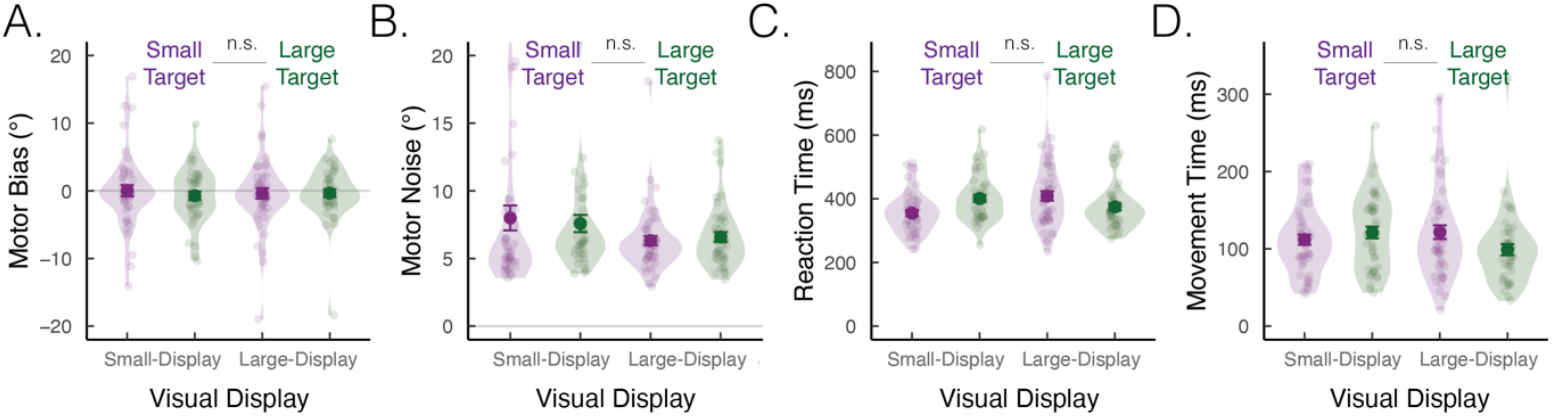
No differences in kinematic features across groups. Groups did not differ in **(A)** motor bias (mean movement angle during baseline), **(B)** motor noise (SD of movement angle during baseline), **(C)** reaction time (from target offset to movement exceeding 1 cm), or **(D)** movement time (from 1 cm movement initiation to target distance at 8 cm). Dots represent individual participants; error bars indicate ±1 SEM around the mean. *n*.*s*. denotes p > 0.05.

## Discussion

We found that goal uncertainty attenuates implicit adaptation, independent of low-level visual or kinematic factors. Controlling for these variables is essential: Enhanced visual salience (e.g., differences in visual feature size) may increase arousal or task engagement, thereby enhancing adaptation (32), while differences in motor kinematics—such as motor noise and movement time—can independently influence learning (33,34). By holding these factors constant, we isolate goal uncertainty as a robust and specific constraint on implicit adaptation.

Our findings reveal that implicit adaptation of reaching movements is modulated not only by sensory uncertainty in a Bayes-optimal manner, but also by uncertainty in goal representation—a factor overlooked in existing models. Traditional computational accounts and empirical tests have focused on how sensory feedback uncertainty governs learning (11,35–37). These models often rely on the Kalman filter, a normative algorithm for updating predictions based on noisy observations. The key parameter is the Kalman gain (*K*), which determines how much the planned motor endpoint (*x*) (in general Kalman terms, the internal state) is adjusted in response to a prediction error (*e*). For perceptually guided reaching, the Kalman gain is typically expressed as:

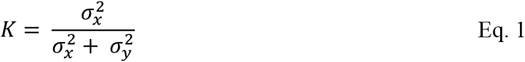

where 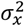 reflects uncertainty in the planned endpoint (which we will assume coincides with the executed movement, for simplicity), and 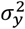 reflects observation noise in sensory feedback about the movement outcome (11). The motor plan is updated trial-by-trial according to:

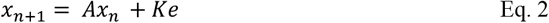

where *A* represents memory retention and *e* is the prediction error, defined as the difference between the observation of sensory feedback about the movement outcome (*y*) and the goal location (ŷ).

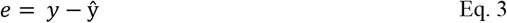

Assuming the sensory feedback about the movement outcome is veridical, but its observation (*y*) is corrupted by visual noise (*εy*), it can be expressed as:

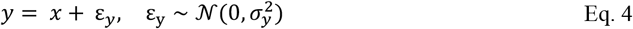

Crucially, this standard formulation implicitly assumes that the goal location (*ŷ*) is known with certainty. Our results challenge this assumption. When the goal is ambiguous—such as when it is spatially extended or poorly defined—this introduces additional uncertainty into the error signal.

To capture this novel idea, the nervous system may jointly represent both the planned movement outcome *and* the goal location:

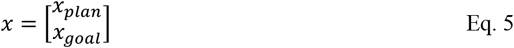

Assuming independent noise in the planned movement outcome 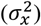 and the sensory feedback about the goal location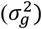, the covariance matrix is:

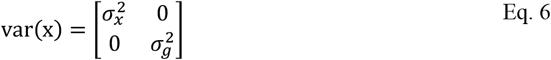

Now, the system observes the discrepancy between the planned movement endpoint and the sensory feedback about the goal location.

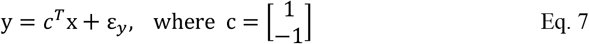

The Kalman gain in this formulation is thus given by:

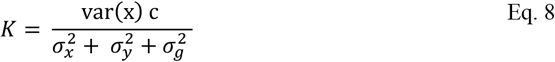

This principled extension reveals that uncertainty in the goal, in terms of the planned endpoint of movement—like uncertainty in sensory feedback about the movement outcome—diminishes the reliability of the error signal and, in turn, attenuates learning. By embedding *both* cursor and goal states within the model, this framework places goal uncertainty on equal computational footing with sensory uncertainty (38). In doing so, it broadens the normative framework of sensorimotor adaptation, offering a more complete account of how the brain regulates learning under uncertainty.

These findings also prompt a re-evaluation of prior studies that manipulated target size to influence implicit adaptation. Previous work showed that when visual feedback landed within a large target (“hit” condition), adaptation was reduced compared to when the same feedback missed a smaller target (“miss” condition) (24,39). This was interpreted as evidence for an intrinsic reward signal—feedback hitting the target was treated as a success, reducing the drive to adapt. However, our results suggest an alternative explanation: the larger target may have introduced greater goal uncertainty, akin to our manipulation using a broader arc. When the goal is spatially diffuse, estimating its midpoint becomes more uncertain, which in turn weakens the error signal and attenuates adaptation—not due to intrinsic reward, but due to degraded goal precision.

These findings also raise new questions about the neural mechanisms by which goal uncertainty modulates implicit adaptation. The dorsal premotor cortex (PMd) has been implicated in representing action goals and encodes multiple potential targets under conditions of uncertainty (31,40,41). Ambiguity in goal representation within PMd may reduce the precision of downstream target signals transmitted to the cerebellum, a structure critical for computing perceptual prediction errors during adaptation (21,42–44). In this framework, degraded goal information would blunt the computation of precise error signals, thereby diminishing cerebellar-dependent learning. Future neurophysiological studies could directly test this hypothesis by examining how uncertainty in PMd target representations alters the fidelity of error signals in the cerebellum—and, in turn, modulates adaptive behavior (45).

## Methods

### Ethics Statement

All participants provided informed consent in accordance with protocols approved by Carnegie Mellon University’s Institutional Review Board. The study was conducted online, and participants received monetary compensation for their time.

### Participants

A total of 200 participants completed the study (Age: 18 – 30 years old). Participants were recruited via Prolific (www.prolific.com) and compensated at $12.00/hour. Eligibility criteria included: (1) ≥95% approval rating, (2) native English speaker, (3) right-handed, and (4) normal or corrected-to-normal vision. The task lasted approximately 30 minutes.

### Sample size

Sample sizes were guided by prior web-based motor adaptation studies (46–49). Each group included 50 participants—substantially larger than typical in-lab studies (∼20 participants/group)—to accommodate potential exclusions due to noisy data (see Exclusion Criteria).

### Apparatus

The experiment was built using OnPoint, a JavaScript-based platform for running customized online motor learning tasks (50) (Figure 1A). Participants completed the task in a web browser using their own devices (Trackpad: 142; Optical Mouse: 37; Trackball: 1). Prior work has shown that neither browser type nor pointing device significantly affects performance (46). Stimulus size and position scaled with each participant’s monitor; for clarity, all parameters reported below reflect values on a standard 13-inch screen.

### Trial Structure

On each trial, participants moved a white cursor (diameter: 0.4 cm) into a central white start ring (diameter: 0.5 cm). Once centered, the ring filled in. After holding for 500 ms, the cursor disappeared, and a blue target appeared along an invisible ring (radius: 8 cm). Targets were presented in a pseudo-random order centered at one of four fixed locations: 45° (upper right), 135° (upper left), 225° (lower left), or 315° (lower right), with one reach to each location per cycle (1 cycle = 4 reaches). The target remained visible for a jittered interval (500–1000 ms) to prevent participants from anticipating its offset, which served as the go cue. Participants were instructed to make a straight reach to bisect the remembered target location. Cursor feedback was displayed during the movement and remained visible for 50 ms upon reaching the target distance. A neutral tone marked movement completion.

### Block Structure

The experiment consisted of three blocks (220 trials total; 55 cycles; Figure 1B): a baseline block with veridical feedback (40 trials; 10 cycles), a perturbation block with rotated feedback (160 trials; 40 cycles), and a no-feedback aftereffect block (20 trials; 5 cycles). In the baseline block, participants were familiarized with the task and instructed: “Your target will be represented as a XX. Slice through the midpoint of the target quickly and accurately,” where XX varied by experimental group. In the perturbation block, visual feedback was rotated by a fixed 15°—clockwise or counterclockwise (counterbalanced across participants). Participants were told: “Ignore the visual feedback; continue to slice through the midpoint of the target quickly and accurately.” In the aftereffect block, no feedback was provided, and instructions reverted to those used in the baseline block.

### Experimental Groups

Participants were randomly assigned to one of four groups in a 2×2 between-subjects design (50/group) (Figure 1C). Goal uncertainty was manipulated by varying the spatial precision with which the target’s midpoint was specified: either a small area (low uncertainty=small-target) or an extended arc (high uncertainty=large-target). To dissociate goal uncertainty from visual format, we independently varied the size of features in the visual display: a sparse dot-based array (small-display) or a continuous illuminated arc (large-display).

This manipulation yielded four experimental groups. In the High-Certainty / Small-Display condition, the target was a single filled circle (diameter = 0.5 cm or 5°). In the High-Certainty / Large-Display condition, the target was a 90° arc (thickness = 0.5 cm) with a central occlusion (0.5 cm), preserving the arc format while highlighting the midpoint. In the Low-Certainty / Small-Display condition, two small arcs (length = 0.5 cm or 5°) marked the endpoints of an implicit 90° arc, requiring participants to mentally interpolate the midpoint from the endpoints. In the Low-Certainty / Large-Display condition, the target was a continuous 90° arc (thickness = 0.5 cm) with no explicit midpoint marker.

### Attention and Instruction Checks

Ensuring participant engagement is a common challenge in online studies. To address this, we included intermittent attention checks within the first 50 trials, prompting participants with simple instructions such as “Press the letter ‘a’ to proceed” or “Press the letter ‘e’ to continue.” To confirm that participants understood the clamped feedback was non-contingent and that movement should begin at target offset, we included a multiple-choice instruction check. Participants who failed two checks were removed from the study.

### Exclusion Criteria

Trials were excluded if the movement angle deviated more than 3 standard deviations from a 10-trial moving average, the movement was directed ≥ 90° away from the target, reaction time exceeded 2000 ms (measured from target offset to movement exceeding 1 cm), or movement time exceeded 1000 ms (measured from movement onset to reaching the target distance). Any participant with more than 10% of trials meeting these criteria was removed from analysis. This resulted in the exclusion of 20 out of 200 participants, yielding a final sample of 180: 46 in the High-Certainty / Small-Display group, 49 in the High-Certainty / Large-Display group, 42 in the Low-Certainty / Small-Display group, and 43 in the Low-Certainty / Large-Display group.

### Primary Dependent Variables

We focused our analyses on the hand position data recorded when the movement amplitude reached the target radius. These data were used to calculate our main dependent variable, movement angle, defined as the difference between the hand position and the target. Movement angles across different perturbation directions were flipped such that positive movement angles always signified performance changes that nullifies the perturbation. Movement angles were also baseline-subtracted to correct for small idiosyncratic movement biases (51,52). Baseline performance included all 10 cycles of the baseline veridical feedback block (trials 1 – 40). Aftereffect performance included all 5 cycles of the aftereffect block (trials 201 – 220).

### Cluster-based Permutation Tests

We used a continuous performance measure to compare the groups, implementing a cluster-based permutation test on the movement angle data (26,53). The test consisted of two steps. First, a F-test (comparing all four experimental conditions) was performed for each movement cycle across experimental conditions to identify clusters that showed a significant difference. Clusters were defined as epochs in which the p-value from the F-tests were less than 0.05 for at least two consecutive cycles. The F values were then summed up across cycles within each cluster, yielding a combined cluster score. Second, to assess the probability of obtaining a cluster of consecutive cycles with significant p-values, we performed a permutation test. Specifically, we generated 1,000 permutations by shuffling the condition labels. For each shuffled permutation, we computed the sum of F-scores to generate a null distribution of scores. The proportion of random permutations which resulted in a F-score that was greater than or equal to that obtained from the data could be directly interpreted as the p-value. Clusters with p_perm_ < 0.05 are reported. Post hoc t-tests were conducted on significant clusters using 1,000 permutations to assess pairwise group differences.

### Model Fitting

To quantify learning dynamics, we fit a standard state-space model to each participant’s trial-by-trial hand angle data during the perturbation block (35,54). This model characterizes how behavior evolves in response to error using three core parameters: the learning rate (K) – also known as the Kalman Gain—reflects the proportion of observed error incorporated into the updated motor plan (x). The retention rate (A) captures how much of the previous plan is carried forward from one trial to the next. A motor noise term 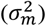 accounts for variability in movement execution unrelated to learning:

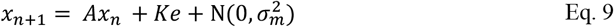

The observed visual error (*e*)—defined as the difference between the observed movement outcome (visual feedback) and the goal location (i.e., the target)—was set to 0° during the baseline block and fixed at 15° during the perturbation block, consistent with the clamped visual rotation. Model fitting was performed using maximum likelihood estimation, implemented in R via the optim function from the stats package.

### Validity of Online Psychophysics

While some may question the validity of online experiments due to variability in input devices such as trackpads or mice, several safeguards support the reliability of our findings. First, our previous work has shown that device type has minimal influence on measures of motor adaptation (46). Second, our group-level comparisons tend to average out individual variability, mitigating the impact of hardware-related noise. Third, we employed rigorous attention and instruction checks, as mentioned above, excluding disengaged participants to ensure data quality. Most importantly, our results replicate hallmark features of implicit adaptation seen in the lab, providing convergent validity that our findings are not artifacts of the online testing environment (50,55).

## Acknowledgements

We thank Matthew Warburton for illustrating our experimental setup.

